# Mapping the Global Network of Extracellular Protease Regulation in *Staphylococcus aureus*

**DOI:** 10.1101/764423

**Authors:** Brittney D. Gimza, Maria I. Larias, Bridget G. Budny, Lindsey N. Shaw

## Abstract

A primary function of the extracellular proteases of *Staphylococcus aureus* is to control the progression of infection by selectively modulating the stability of virulence factors. Consequently, a regulatory network exists to titrate protease abundance/activity, to influence accumulation, or lack thereof, of individual virulence factors. Herein, we comprehensively map this system, exploring regulation of the four protease loci by known and novel factors. In so doing, we determine that seven major elements (SarS, SarR, Rot, MgrA, CodY, SaeR, and SarA) form the primary network of control, with the latter three being the most powerful. We note that expression of aureolysin is largely repressed by these factors, whilst the *spl* operon is subject to the strongest upregulation of any protease loci, particularly by SarR and SaeR. Furthermore, when exploring *scpA* expression, we find it to be profoundly influenced in opposing fashions by SarA (repressor) and SarR (activator). We also present the screening of >100 regulator mutants of *S. aureus*, identifying 7 additional factors (ArgR2, AtlR, MntR, Rex, XdrA, Rbf, and SarU) that form a secondary circuit of protease control. Primarily these elements serve as activators, although we reveal XdrA as a new repressor of protease expression. With the exception or ArgR2, each of the new effectors appear to work through the primary network of regulation to influence protease production. Collectively, we present a comprehensive regulatory circuit that emphasizes the complexity of protease regulation and suggest that its existence speaks to the importance of these enzymes to *S. aureus* physiology and pathogenic potential.

**Importance:** The complex regulatory role of the proteases necessitates very tight coordination and control of their expression. Whilst this process has been well studied, a major oversight has been the consideration of proteases as a single entity, rather than 10 enzymes produced from four different promoters. As such, in this study we comprehensively characterized the regulation of each protease promoter, discovering vast differences in the way each protease operon is controlled. Additionally, we broaden the picture of protease regulation using a global screen to identify novel loci controlling protease activity, uncovering a cadre of new effectors of protease expression. The impact of these elements on the activity of proteases and known regulators was characterized producing a comprehensive regulatory circuit that emphasizes the complexity of protease regulation in *Staphylococcus aureus*.

## Introduction

*Staphylococcus aureus* is an opportunistic human pathogen known for causing both hospital- and community-acquired infections. It is capable of causing a plethora of diseases that range from minor skin and soft tissue infections, such as boils and carbuncles, to septicemia, endocarditis, osteomyelitis and toxic shock syndrome [1–3]. This broad disease potential can be attributed to the coordinated production of a wealth of virulence factors by *S. aureus* within the human host. Collectively these elements allow the pathogen to evade phagocytosis, promote abscess formation, travel from initial sites of infection to invade new tissues, and to induce a variety of syndromes [4]. These virulence causing entities can be divided into two broad groups: adherence factors and exoproteins. Adherence factors are responsible for the attachment of *S. aureus* to host tissues so that colonization may occur [5], and can also interfere with the host immune system to facilitate immune evasion [6]. Conversely, exoproteins are secreted by *S. aureus* and function to acquire nutrients by breaking down host tissues and, more importantly, target the immune system, engendering immune subversion [7].

As part of this cadre of secreted factors are 10 extracellular proteases, which are produced by almost every *S. aureus* strain (**Fig. 1**) [8, 9]. These include: a metalloprotease, aureolysin (*aur*); a serine protease V8 (*sspA*); two cysteine proteases, staphopain B (*sspB*) and staphopain A (*scpA*); and six serine-protease-like enzymes (*splABCDEF*) [9, 10]. The functions of these enzymes have been studied by ourselves and others, and includes their ability to hydrolyze a variety of host proteins, as well as self-derived toxins. With regards to host factors, the secreted proteases have been demonstrated to proteolyze proteins such as fibrinogen, elastin, and the heavy chains of immunoglobulins to promote tissue invasion, immune system evasion, and the dissemination of infection [11–13]. In the context of the self-degradome, these enzymes can cleave multiple virulence determinants to promote bacterial invasion, immune evasion, and survival. For example, Aureolysin was shown to control the stability of both phenol soluble modulins and α-toxin [14, 15], as well as the adhesin clumping factor B (ClfB) [16]; whilst SspA is able to cleave surface protein A (SpA) and the fibrinogen-binding proteins (FnBPs) [17, 18].

**Figure 1.**
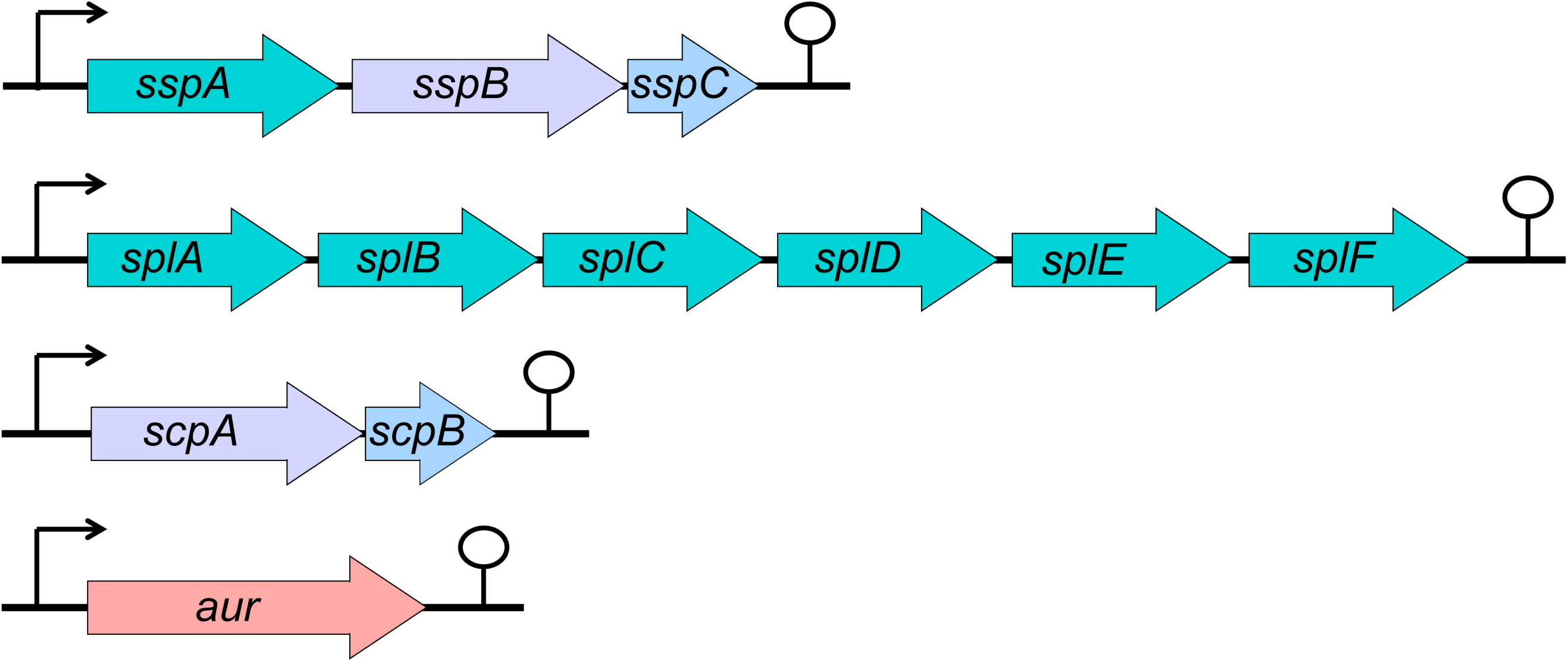
Genetic Organization of the *S. aureus* Secreted Proteases Loci. The colors of arrows are representative of catalytic activity classification: metalloprotease in pink; serine proteases in green; cysteine proteases in purple; and the inhibitors of the staphopains (the staphostatins) in blue.

Recently, our group assessed the importance of secreted proteases in *S. aureus* pathogenesis using a strain where all 10 enzymes were deleted [19]. Here, we demonstrated that secreted proteases are required for growth in whole human blood, serum, peptide-rich media, and in the presence of antimicrobial peptides. Additionally, these enzymes are also necessary for *S. aureus* to resist phagocytosis by human granulocytes and monocytes. Most striking, however, were the *in vivo* phenotypes of this mutant, where a decrease in dissemination and abscess formation was observed in infected mice compared to the wild-type. Conversely, when assessing mortality, the complete protease-null strain demonstrated pronounced hypervirulence. These contrasting phenotypes were explained using proteomics, where an increase in stability of secreted and surface-associated virulence factors was demonstrated en masse in the mutant, thus facilitating more aggressive and deadly infections. Importantly, many of these findings were also demonstrated in a companion study by Zielinska et al [20]. As such, it would appear that secreted proteases have a biphasic role in infection, serving on the one hand to modulate the stability of self-derived pathogenic determinants, so as to control disease severity and progression, whilst at the same time playing their own direct role by cleaving host proteins to promote invasion, immune evasion, and survival.

Given the complex regulatory role of *S. aureus* proteases during infection, it follows that there must be, and indeed is, tight control of their expression mediated by a collection of different factors. This is evidenced by the number of elements that have been identified thus far as influencing protease production, including RNAIII SarS, SarR, SarA, SarV, SarX, SarZ, ArlRS, CodY, Rot, MgrA, and SaeRS [21–33]. Of these factors, SarS, SarR, CodY, Rot, MgrA, SaeR and SarA are considered the primary regulators, with each being shown to directly influence protease transcription [21–27]. A major oversight when studying the control of protease production in *S. aureus*, however, has been the consideration of these factors as a single entity, rather than 10 enzymes produced from four different promoters. Of the seven major regulators, only SarA and Rot have been explored in the context of all four protease promoters [9, 10], with SarA shown to specifically repress the transcription of *aur*, *scpA*, *ssp* but not *spl* [9, 10]; whilst Rot has been described as a direct negative regulator of all secreted protease operons [23]. For the other primary regulators, CodY has been shown to directly repress *ssp* transcription [22], whilst SarS and SarR have been explored only in the context of *aur* and *ssp* promoter binding [21]. Finally, MgrA has been shown to activate *aur*, *ssp*, and *spl* transcription [25, 34], whilst SaeR has been described as an activator for *spl,* but a repressor for *aur* [24].

Consequently, the overarching goal of this study was to explore and further our understanding of the regulation of secreted proteases by known regulatory factors in *S. aureus*, whilst concurrently uncovering new effectors of protease transcription. Accordingly, we present a comprehensive mapping of protease regulation by all known *S. aureus* transcription factors in CA-MRSA strain USA300.

## Materials and Methods

### Media and growth conditions

All cultures were grown overnight at 37°C with shaking at 250 rpm in 5 mL of either tryptic soy broth (TSB) or lysogeny broth (LB). When required, antibiotics were added at the following concentrations: *E. coli* - 100 μg ml^−1^ ampicillin, 12.5 μg ml^−1^ tetracycline; *S. aureus* - 5 μg ml^−1^ tetracycline, 5 μg ml^−1^ erythromycin, 25 μg ml^−1^ lincomycin, and 2.5 μg ml^−1^ chloramphenicol. To obtain synchronous cultures, overnight *S. aureus* cultures were diluted 1:100 into 5 mL of fresh media and grown for 3h, before being standardized to an optical density (OD_600_) of 0.05 in 100 mL of fresh TSB. When assessing growth, OD_600_ was measured hourly using a Synergy 2 plate reader (Bio-Tek).

### Bacterial strains

All bacterial strains and plasmids used in this study are listed in **Table 1**. Transposon mutants for all available transcriptional regulators in *S. aureus* USA300 JE2 were obtained from the Nebraska Transposon Mutant Library (NTML). Those subjected to further study were transduced into USA300 Houston, as described by us previously [35], using phi11. The construction of an *mgrA* mutant in *S. aureus* Becker was previously described [36]. This mutation was transduced into USA300 Houston using phi85.

**Table 1.** Strains and plasmids used in this study.

| Strain or plasmid | Description* | Reference or source |
| --- | --- | --- |
| *E. coli* |  |  |
| DH5α | Cloning strain | [1] |
| *S. aureus* |  |  |
| RN4220 | Restriction-deficient strain | Lab stock |
| USA300 HOU | USA300-HOU MRSA Isolate | [2] |
| BDG2625 | USA300 HOU *codY*::Tn::erm *codY*<sup>-</sup> | This study |
| BDG2623 | USA300 HOU *sarS*::Tn::erm *sarS*<sup>-</sup> | This study |
| BDG2621 | USA300 HOU *sarA*::Tn::erm *sarA*<sup>-</sup> | This study |
| BDG2624 | USA300 HOU *rot*::Tn::erm *rot*<sup>-</sup> | This study |
| BDG2622 | USA300 HOU *saeR*::Tn::erm *saeR*<sup>-</sup> | This study |
| CYL1040 | Becker *mgrA*::cm *mgrA*<sup>-</sup> | [3] |
| BDG2626 | USA300 HOU *mgrA*::cm *mgrA*<sup>-</sup> | This study |
| BDG2479 | USA300 HOU *sarR*::tet *sarR*<sup>-</sup> | This study |
| BDG2331 | USA300 HOU *sarU*::Tn::erm *sarU*<sup>-</sup> | This study |
| BDG2333 | USA300 HOU *rex*::Tn::erm *rex*<sup>-</sup> | This study |
| BDG2329 | USA300 HOU *rbf*::Tn::erm *rbf*<sup>-</sup> | This study |
| BDG2334 | USA300 HOU *argR2*::Tn::erm *argR2*<sup>-</sup> | This study |
| BDG2336 | USA300 HOU *atlR*::Tn::erm *atlR*<sup>-</sup> | This study |
| BDG2328 | USA300 HOU *mntR*::Tn::erm *mntR*<sup>-</sup> | This study |
| BDG2332 | USA300 HOU *xdrA*::Tn::erm *xdrA*<sup>-</sup> | This study |
| LES55 | SH1000 *sigS*::tet *sigS*<sup>-</sup> | [4] |
| Plasmids |  |  |
| pJB38 | Plasmid to create mutants in *S. aureus* | [5] |
| pBDG01 | pJB38 construct for *sarR* mutation, Amp<sup>R</sup> CM<sup>R</sup> | This study |

\*The following abbreviations specify resistance cassettes to: Erm – erythromycin, CM – chloramphenicol, Tet – tetracycline, Amp – ampicillin.

### Construction of a *sarR* mutant strain

A tetracycline marked disruption of *sarR* was generated using pJB38, as described by Bose et al. [37]. Regions up and downstream of *sarR*, including portions of the 5’ and 3’ ends of the coding gene, were amplified via PCR using primers OL4208/OL4209 and OL4210/OL4211. A tetracycline resistance cassette was amplified using OL4299/OL4300 from a SH1000 *sigS*::*tet* mutant [38]. Using MluI sites, the tetracycline cassette was ligated between the upstream and downstream fragments of *sarR*, and ligated directly into pJB38 using EcoRI and KpnI sites. Using the established protocol, the majority of *sarR* was deleted in USA300 Houston using allelic replacement [37]. Strains were confirmed by PCR and sequencing (Eurofins Genomics) using primers OL4577/OL4578, which amplify across the deleted region where the tetracycline cassette was inserted.

### Quantitative Real-Time PCR Analysis

To quantify expression changes for target genes (Primers are listed in **Table S1**), quantitative real-time PCR (qRT-PCR) was performed, as described by us previously [39]. All targets were normalized using 16s rRNA expression and fold change from the wild-type was determined using the 2^−ΔΔC^_T_ method [40].

### Zymography

Strains grown for 15h overnight were adjusted to equal optical densities and pelleted. When assessing proteolytic activity over time, synchronized cultures were grown to exponential phase and standardized to OD_600_ of 0.05 in 100mL of TSB. At the desired time points, cells were pelleted. Thereafter, for all samples, 2 mL of supernatant was processed through an Amicon Ultra 3K centrifugal filter for 60 minutes at 4,000 x g. Concentrated supernatants were recovered by removing filtrate collection tubes, inverting filter devices and spinning again for 2 minutes at 1,000 x g. An equal volume of Laemmli loading buffer was added to the concentrated supernatants and incubated for 30 minutes at 37°C. Next, 20 μl of each sample was loaded onto pre-prepared SDS-PAGE gels containing 0.1 % gelatin, and run until the dye front reached the edge of the plates. Gels were washed twice using 2.5% Triton X-100 at room temperature. Following a rinse with diH_2_O, developing buffer (0.2M Tris, 5mM CaCl_2_, 1mM DTT, pH 7.6) was added and gels were incubated overnight at 37°C static. After incubation, gels were rinsed with diH_2_O and covered with 0.1% amido black for 1h. Once gels were stained, destain 1 (30% Methanol, 10% Acetic Acid) was added for 5-10 minutes, replaced with destain 2 (10% Acetic Acid) until bands became clear, and then replaced with destain 3 (1% Acetic Acid) for storage. Changes in band intensity were quantified using the ImageJ software.

## Results and Discussion

### Exploring the Differential Regulation of Protease Expression by Primary Regulators

To date, seven different transcriptional regulators (Rot, CodY, SarA, SarS, MgrA, SarR and SaeR) [21–27] have been identified as being the primary modulators of secreted protease expression. An oversight, however, is the consideration of *S. aureus* proteases as a single entity, rather than 10 enzymes produced from four distinct loci (**Fig. 1**). Thus, although these elements do indeed have the capacity to regulate the expression of one or more protease, only a handful have been explored in the context of all four operons. Therefore, our initial goal was to fill in missing gaps using qRT-PCR. To assess this, wild-type and regulator mutant strains were grown to post-exponential phase (5h), which is a known window of peak protease expression [9], and assessed for the expression of each protease operon.

We began with the best studied regulator, SarA, whose ability to repress the transcription of *aur*, *scpA*, *ssp* but not *spl* has been previously well established [9, 10]. Here, our analysis provided the expected results: in the absence of SarA, there was a 275-fold increase in *aur*, 10.9 in *scpA*, 23.7 in *sspA* transcript levels, alongside no changes in *spl* expression (**Fig. 2A**).

**Figure 2.**
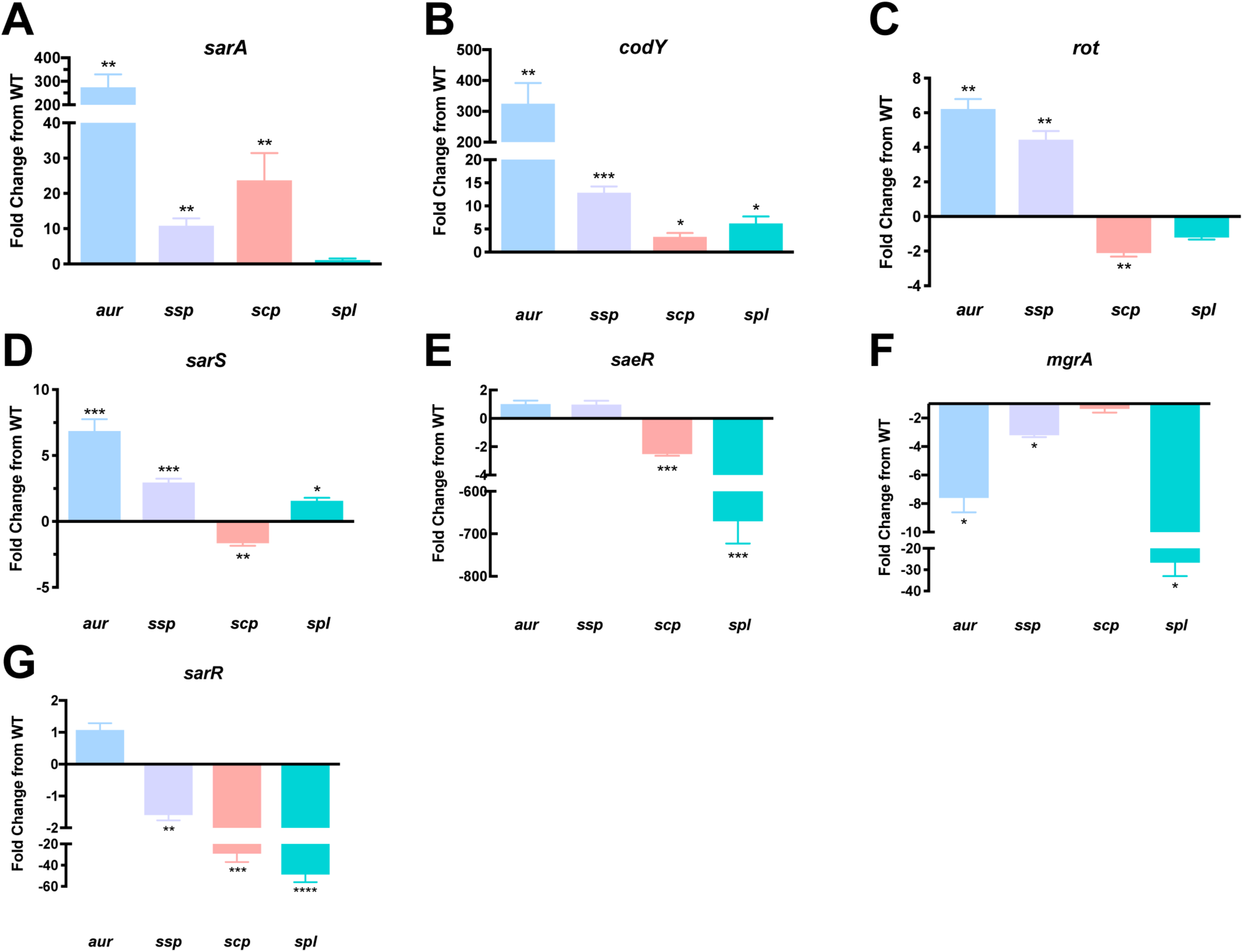
Individual Protease Loci are Differentially Controlled by Major Regulators of *S. aureus*. qRT-PCR was used to determine transcript levels for *aur*, *scp*, *ssp* and *spl* in regulator mutants after 5h of growth. The strains used were: WT = USA300 HOU, alongside mutants for *sarA*, *codY*, *rot*, *sarS*, *saeR*, *mgrA*, and *sarR*. RNA was isolated from three independent cultures. The 16s rRNA gene was used as an internal control. Fold change from WT was determined using the 2^−ΔΔC^_T_ method. A Student’s t-test was used to determine statistical significance, *=p< 0.05, **=p< 0.01, ***=p<0.001, ****=p<0.0001. Error bars shown ± SEM.

Next, we investigated CodY, whose ability to influence protease expression was identified by microarray analysis in UAMS-1 [22]. Here, Majerczyk et al. found that in the absence of CodY, *sspA* had increased transcript levels. Additionally, in the same study, CodY was shown to bind the *spl*, *sspA*, and *aur* promoters, however the binding to *aur* and *spl* was deemed biologically irrelevant as changes in their expression were not observed upon *codY* deletion. As such, the ability of CodY to modulate expression of *aur*, *scpA,* and *spl* has not been previously described. Herein, in the absence of CodY we observed a significant 324-fold increase in *aur*, 12.8 in *sspA*, 3.3 in *scpA* and 6.2 in *spl* transcript levels (**Fig. 2B**). Collectively this data suggests that CodY is a negative regulator of secreted protease expression that rivals SarA in its potency.

We next considered Rot, which was first shown to negatively regulate *sspA* and *spl* transcription in a RN6390 microarray [41]. In another study assessing *aur* and *sspA* regulation in 8325-4, Rot functioned as a direct repressor of both loci [25]. In support of these studies, others have demonstrated that Rot represses *aur* and *sspA*, whilst also directly repressing *spl* through promoter binding in LAC [23]. Additionally, in the same study Rot was shown for the first time to directly repress *scpA* transcription. In our study, upon *rot* inactivation, there was a significant fold increase of 6.2 for *aur* and 4.5 for *sspA* transcript levels, which is in line with previous research [23]. Additionally, a significant 2.1-fold decrease in *scpA* expression alongside no change for *spl* was observed, contradicting previous studies, where increased transcription for both was observed upon *rot* deletion (**Fig. 2C**). We note, however, that previous studies regarding Rot regulation differ from ours through the use of media supplemented with different nutrients. Specifically, in work by Mootz et al. growth media was supplemented with glucose, which has been documented as repressing the *agr*-quorum sensing system via the decreased pH produced from carbon metabolism [42, 43]. As such, this decrease in *agr* activity could alter the expression of downstream factors also capable of regulating the secreted proteases. Similarly, Said-Salim et al. used casamino acids-yeast extract—glycerolphosphate broth for their studies. Here, the addition of glycerol, as well as the use of an entirely different complex media, would alter the activity of other transcriptional regulators such as CodY, CcpE, CcpA and RpiRC, which are known to sense the carbon status of the cell [44]. Therefore, whilst Rot has the potential to regulate all four protease loci, our data suggests that Rot primarily controls expression of *aur* and *sspAB*, likely in an *agr-*dependent manner.

SarS was formerly shown to have no significant effect on *aur* and *ssp* transcription during investigation in 8325-4 [27]. Oscarsson et al. however, established that when *sarS* is overexpressed in 8325-4, *aur* and *sspA* transcription is suppressed [25]. In support of a role in *sspA* regulation, another study showed that SarS could bind the *sspA* promoter [27]. To date, the effects of SarS on *scpA* and *spl* transcription have not yet been investigated. Our analysis of protease transcription in the absence of SarS revealed a significant increase for *aur* (6.9-fold), *sspA* (2.9-fold) and *spl* (1.6-fold), but a 1.7-fold decrease in *scpA* transcript levels (**Fig. 2D**). This data thus supports a role for SarS as a repressor of *aur* and *sspA* expression, and identifies the *spl* operon as a new target of negative regulation by this factor. Conversely, we reveal *scpA* as a being activated by SarS, demonstrating, as with our data for Rot, that each of the four proteases are often subject to differential and opposing regulation by the same element.

The ability of SaeR to influence protease expression was previously described by microarray analysis, where, in the absence of SaeR/S in LAC, there was a decrease in *spl* transcription [24]. Furthermore, in this same study it was observed that this effect was direct, as SaeR was shown to bind to the *spl* promoter. Additionally, in the same background Cassat et al., showed a decrease in SplA*-*F protein levels following *sae* inactivation [45]. In support of this, we observed a striking 671-fold decrease in *spl* transcript levels upon *saeR* deletion, which is the most pronounced alteration in expression for any protease observed in this study (**Fig. 2E**). With regards to *aur*, the previously referenced studies revealed an increase in *aur* transcription [24], as well as an increase in Aur protein levels [45] in the absence of *saeRS*. In our study, however, no change in transcription was observed, which is in line with Oscarsson et al., who derived similar findings in RN6390 [25]. Of note, the changes observed during microarray and proteomic analysis were performed during stationary phase, rather than post-exponential phase. Therefore, the disagreement regarding *aur* regulation could be a product of different time points used for assessment. This is supported by our observation that, when analyzed throughout growth, SaeRS is the only major regulator in *S. aureus* to demonstrate a rebound in transcriptional activity during stationary phase (our unpublished observation). This suggests that SaeRS may have varying or biphasic functions with regards to virulence factor regulation during *S. aureus* growth. Regarding *scpA*, the effect of SaeR on transcription has not until now been investigated. Herein, we observed a 2.5-fold decrease in *scpA* transcription in the absence of SaeR, indicating that, like the *spl*s, it is activated by this factor. Lastly, no change in *sspA* transcription was observed, which, whilst in line with Oscarsson et al [25], contradicts Cassat et al [45], who observed an increase in SspA and SspB protein levels during stationary phase. As previously suggested, this conflict is likely explained by the varying impact of SaeRS during different growth phases. As such, our data supports a role for SaeR during post exponential growth in the activation of *spl* and identifies *scpA* as a new target for SaeR upregulation.

We next investigated MgrA, which, has previously been shown to activate *aur* and *sspA* transcription in 8325-4 [25]. Using RNA-sequencing in LAC, others have shown that the absence of MgrA produced decreased *aur* and *spl* transcript levels [34]. Herein, in correlation with previous studies, we observed a significant 7.6-fold decrease in *aur*-, 3.2 in *sspA*-, and 26.7 in *spl*-transcript levels (**Fig. 2F**). Lastly, until now, the effect of MgrA on *scpA* had not been investigated. In our study, no changes in *scpA* transcript levels were identified, which again demonstrates differential regulation of the various protease loci. This is particularly interesting, as it is an additional example of the two staphopain enzymes (SspB and ScpA), which share strong homology [46–48], although quite different substrate specificities [48], as being regulated in opposing fashions.

Finally, we investigated SarR, which has formerly been shown to positively affect *aur* and *sspA* transcript levels in 8325-4 [21]. In contrast, in another study it was shown to negatively affect *aur* when overexpressed in an 8325-4 *agr/sarA* double mutant [25]. Interestingly, however, in our study, no change in *aur* transcript levels were detected in the absence of *sarR*. When considering *ssp* expression, we did observe a significant 1.6-fold decrease in transcript levels (**Fig. 2G**) in the *sarR* mutant, which is in correlation with Gustafsson et al.. With regards to *scpA* and *spl*, SarR has not been previously investigated as controlling their transcription. Herein, we observed a significant 29.1-fold decrease in *scpA* and 48.8-fold decrease in *spl* transcript levels. Our data therefore supports a role for SarR in upregulating the *ssp* operon to a minor extent, whilst serving as one of the strongest activators of *scpA* and *spl* expression identified thus far.

### Defining the Pathway of Control for Secreted Protease Expression by Known Major Regulators

Collectively, our findings confirm 14 regulatory pathways for secreted protease transcription, whilst at the same time identifying eight new nodes of expression (**Fig. 3**). For *aur*, we found it to be regulated by CodY in addition to SarA, Rot, SarS and MgrA. Interestingly, with the exception of MgrA, each of these factors engenders repression of *aur* expression, with some (SarA and CodY) exerting profound influence. This is perhaps explained by the observation that Aureolysin sits atop the protease activation cascade, which flows from Aur to V8 and then Staphopain B [11, 49–51]. As such, repressing Aureolysin would allow the *S. aureus* cell to keep the majority of proteases activity restrained by the single act of limiting expression from P*_aur_*. This would be to the cells advantage as, although proteases are undoubtedly valuable enzymes with important roles, they are also at heart destructive in nature. Thus, limiting their activity until it is absolutely required is a major goal with living cells from all kingdoms [52, 53]. This would be particularly true of Aureolysin, given that it has amongst the broadest substrate specificities of any *S. aureus* protease [54]. In the context of enzymes from the *ssp* operon, we did not identify new regulatory nodes, but confirmed their broad regulation, albeit at modest levels, in a fashion that closely resembles that of *aur* control. This finding is again logical, given that the enzymes produced from these loci are part of the protease activation cascade referenced above.

**Figure 3.**
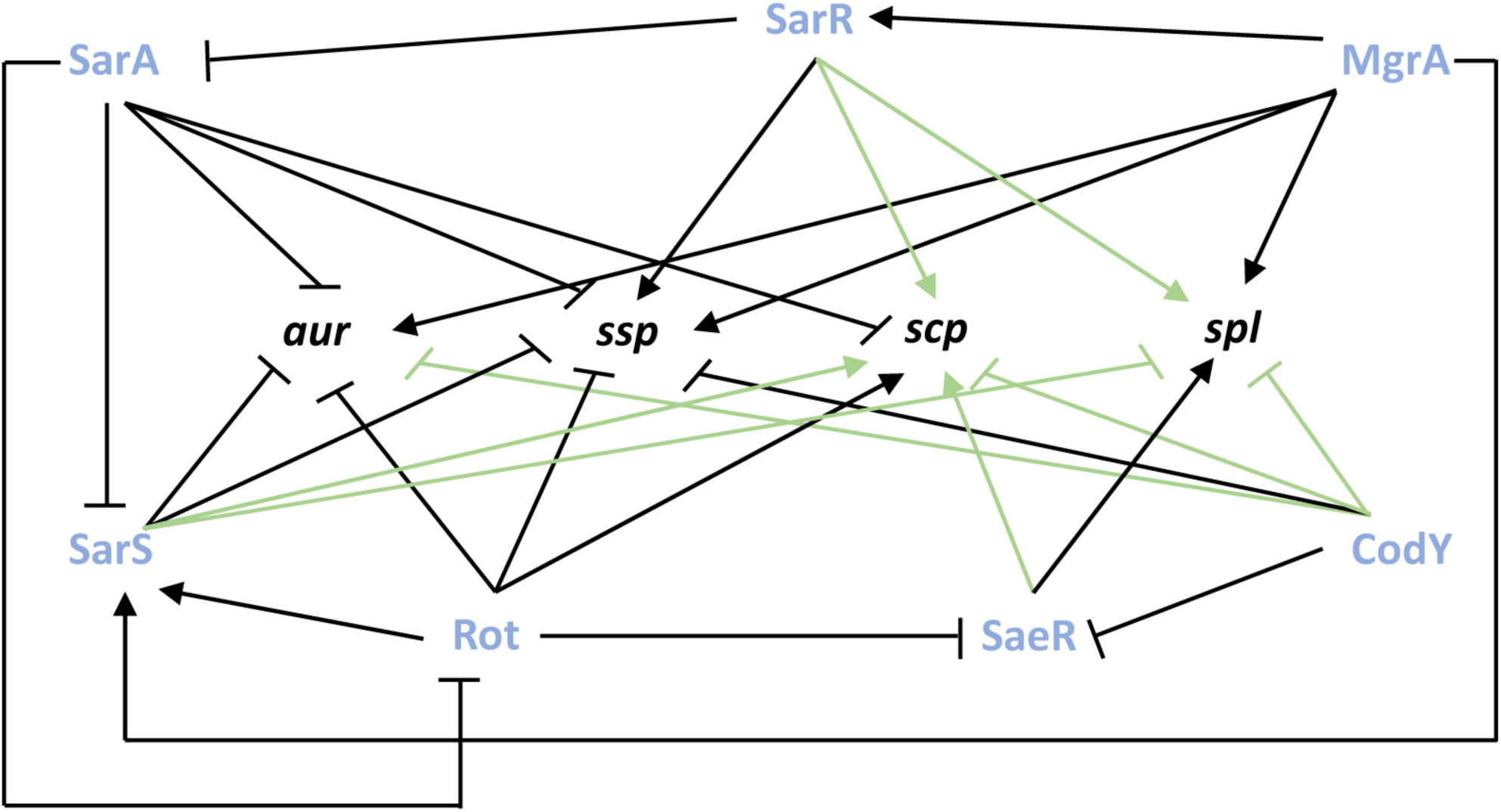
The Primary Network of Control for Individual Protease Loci. Shown are transcriptional regulation events for the seven primary protease regulators of *S. aureus* on the four individual protease loci. Bars indicate repression and arrows activation. New regulatory pathways identified herein between the primary regulators and the protease loci are shown in green.

Interestingly, much of the new knowledge generated herein involves the regulation of the more underappreciated proteases, Staphopain A and the Spls. While the importance of *scpA* in virulence has been shown through *in vivo* studies, as well as by its ability to cleave specific host proteins [13, 55, 56], its transcriptional regulation has been underexplored. While it has been shown previously that *scpA* is regulated by Rot and SarA, we identified herein that SarS, CodY, SaeR, and SarR also control its expression. Whilst much of this regulation is at modest levels, *scpA* expression is profoundly influenced in opposing fashions by SarA (repressor) and SarR (activator). This presents a scenario whereby the presence of this enzyme during infection could be discretely titrated, with high SarA activity resulting in decreased Staphopain A, whilst elevated SarR levels would engender significant production of this enzyme. This could then provide rapid niche specific control of the pathogenic process through Staphopain A activity (or lack thereof) towards self- and host derived proteins. The need for such a network of opposing and stringent control is supported by the observation that Staphopain A is one of only two *S. aureus* secreted proteases with a broad and promiscuous substrate specificity (Aureolysin being the other) [57]; thus, tightly modulating its influence is a necessity for a coordinated and controlled infectious process.

When exploring control of *spl* expression, we note that this operon is subject to some of the strongest regulation observed for any protease loci in this study. Specifically, MgrA, SarR and SaeR each bring about profound upregulation of the *spl* operon, to levels that rival and, in the case of SaeR, exceed that of SarA and CodY for protease control. This is of interest because the Spls are well known for their narrow substrate specificity [58–60]. Indeed, these enzymes share strong homology and many enzymatic characteristics with the exfoliative toxins of *S. aureus*. In the case of these latter proteases, they have only a single known target – desmoglein-1 in the skin of humans, the cleavage of which results in Scalded Skin Syndrome [61]. The Spl enzymes are projected to have a similarly narrow range of substrates [62], thus it is logical that the cell would limit the production of these enzymes until it finds itself in an environment where their activity would prove beneficial. As such, it is logical that the presence and activity of the Spl enzymes can be selectively and rapidly stimulated by these regulatory factors in response to environmental cues within the host, to facilitate infection.

### The Identification of a Cadre of New Effectors of Protease Activity

Given the complex regulatory function of *S. aureus* secreted proteases, tight modulation of their expression is required. As such we set out to more deeply characterize their network of control by uncovering novel effectors of their activity. This was achieved by screening all 108 available transcriptional regulator mutants within the NTML [63] for alterations in proteolytic capacity. Culture supernatants from all strains grown for 15h (a window of peak accumulation for secreted proteases) were prepared and subjected to zymography using gelatin as a substrate, as described by us previously [9]. Of the 108 mutants screened, five of the seven primary regulators (*sarS*, *saeR*, *rot*, *sarA* and *codY*) were included as controls (*sarR* and *mgrA* mutants are not present in the NTML), alongside two other major regulators of protease production: *agrA* and *sigB*. As expected, an increase in proteolytic activity was observed with *sarS*, *rot*, *sarA*, *codY* and *sigB* mutants, whilst a decrease was observed for *saeR* and *agrA* mutants, in comparison to wild-type (**Fig. 4**). For all strains, the intensities of proteolytic banding resulting from gelatin degradation was assessed visually and by densitometry using Image J software (**Fig. 5**).

**Figure 4.**
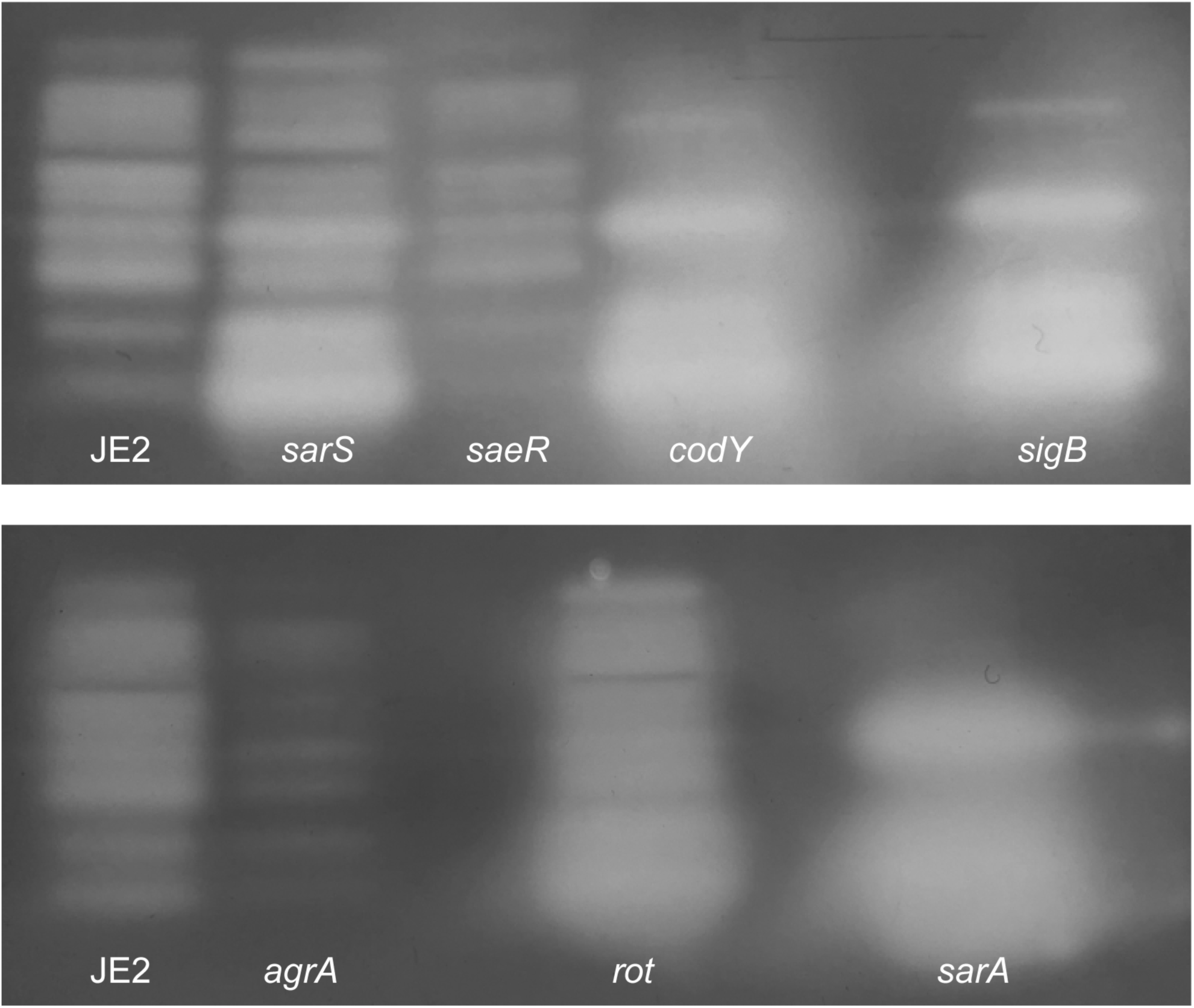
The Impact of Primary Regulator Mutation on Protease Activity. Gelatin zymography was performed to visualize protease activity on 15h culture supernatants obtained from USA300 JE2 and mutants of *sarS*, *saeR*, *codY*, *sigB*, *agrA*, *rot*, and *sarA*.

**Figure 5.**
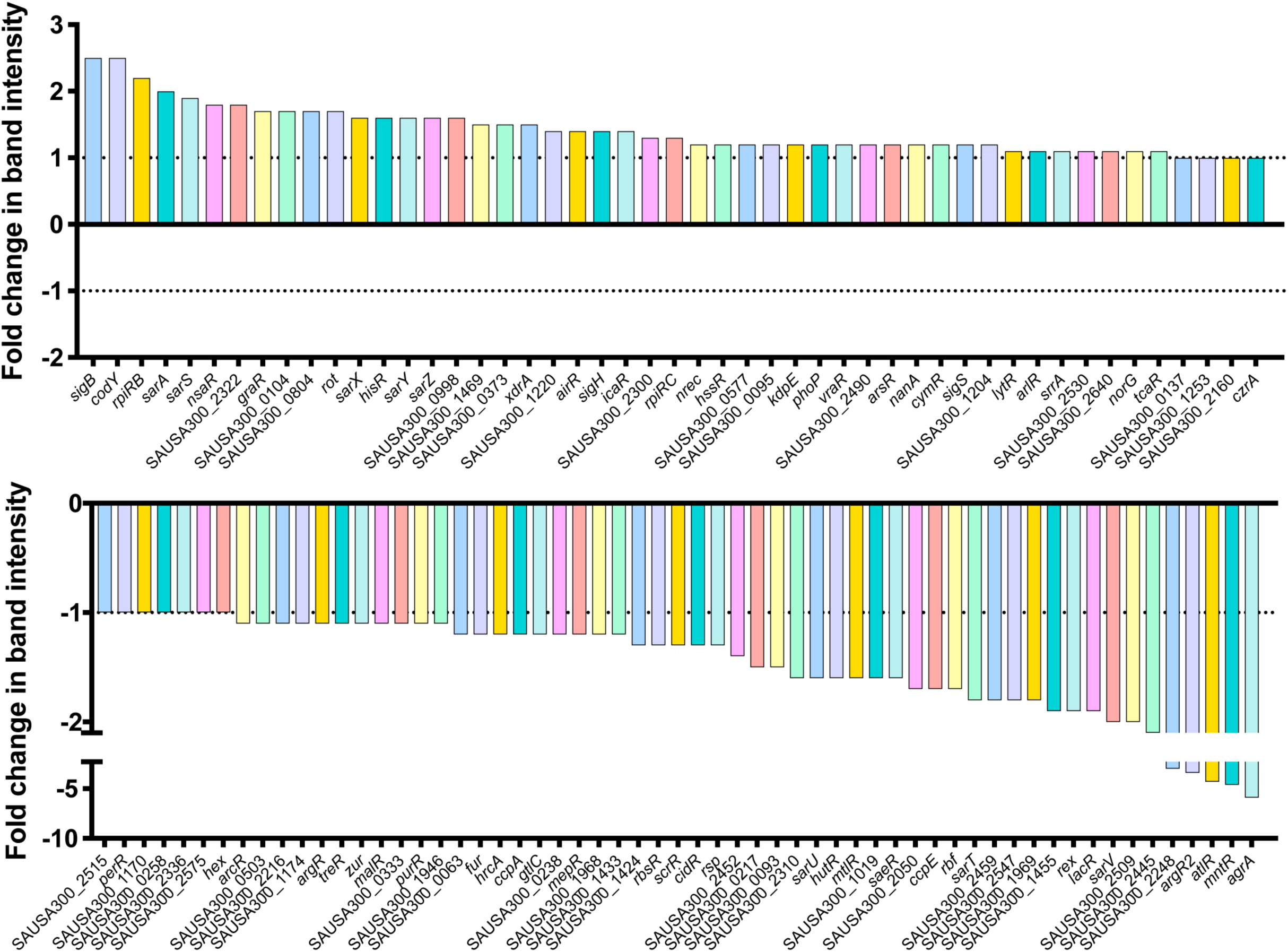
Quantitative Profiling of Protease Activity for All Available Regulator Mutants of *S. aureus*. Zymogram band intensities from all 108 regulator mutants contained within the NTML were measured using densitometry (Image J). Depicted is fold change of band intensity relative to the USA300 JE2 wild-type strain.

Excluding the known major regulators, a total of 45 mutants were identified as having notable alterations in proteolytic activity from our screen, with twenty-six found to have decreased proteolysis (**Table S2**), whilst 19 had an increase (**Table S3**). When assessing mutants that showed increased proteolysis, we identified SarX and NsaR, which have both been previously identified as regulating proteases. SarX has been shown to repress *sspA* transcription in RN6390 [31], whilst NsaR, was shown to be a repressor of *scpA*, *sspA* and *splA-F* in SH1000 [64]. When considering mutants that had decreased proteolysis, we noted SarV and CcpE, both of which have been implicated in modulating protease activity. Specifically, *sarV* disruption in RN6390 led to a decrease in transcription for *aur*, *scpA*, and *splA* [32], whilst loss of *ccpE* in Newman results in impaired expression of all protease loci [65].

Beyond these known factors, we identified a number of intriguing regulators which have yet to be implicated in protease regulation. Of these, several displayed a prominent decrease in protease activity, including SarU. This regulator is an understudied transcription factor belonging to the Sar family, with many of its counterparts already being known to have a role in regulating protease production [66]. In addition, a notable decrease in protease activity was also observed for *rbf* and *atlR* mutants, which encode regulators known to control biofilm formation [67–69]. Further, Rex and MntR, both of which regulate different aspects of cellular metabolism, also caused pronounced decreases in protease activity upon ablation. We also observed a decrease in protease activity upon disruption of *argR2*, which is located within the arginine catabolism metabolic element (ACME) found in USA300 strains [70]. Finally, XdrA/*xdrA*, which has a role in immune evasion via its involvement in the production of protein A [71], was found to produce a notable increase in protease activity upon disruption.

### Exploring Protease Control via a Secondary Network of Regulation

To more deeply explore the new protease regulatory factors identified herein, the seven referenced above were chosen for more detailed study. First, each mutation was transduced into a clean USA300 HOU background and protease activity was continuously throughout growth (**Fig. S1**). In agreement with results from our zymography screen, a decrease in protease activity was observed at all time points for mutants of: *argR2*, *mntR*, *atlR*, *rbf*, *sarU* and *rex*; whilst the *xdrA* mutant demonstrated a minor decrease in protease activity at early times points, but produced the expected increase in proteolysis thereafter. In order to assure that the changes observed were not the result of a simple growth defect, growth curves were performed with all strains, revealing no notable alterations when compared to the wild-type (**Fig. S2**).

Our next step was to determine if the changes observed in the novel regulatory mutants were driven by changes at the level of transcription. Thus, qRT-PCR analysis for each protease loci was performed for the wild-type and regulator mutant strains during post-exponential phase, with the exception of *argR2*, which appears to most notably alter proteolysis at 3h of growth; thus, this time point was used for this strain. When studying changes in the *argR2* mutant, a 1.6-fold decrease in *aur*, 1.8 in *sspA*, and 1.7 in *spl* transcripts were observed (**Fig. 6A**), alongside no change in *scpA* transcription. Next, with *atlR*, we observed a significant 2-fold decrease in *aur* and a 2.2 in *spl* transcripts (**Fig. 6B**), whereas with *scpA* and *sspA*, no changes in transcript levels were noted. For *mntR*, we observed a significant 2.5-fold decrease in *sspA* and 1.7-fold in *spl* transcript levels (**Fig. 6C**), with no changes detected for *aur* and *scpA*. In the context of *rex,* a significant 3.3-fold decrease was seen with *sspA* transcript levels, whilst there were no changes in transcription for the other protease loci (**Fig. 6D**). Following this, we investigated *xdrA* where we observed a significant 1.9-fold increase for *aur* and 4.2-fold in *scpA* transcript levels (**Fig. 6E**), however, with *spl* we observed a significant 2.4-fold decrease in expression. When studying the *rbf* mutant there was a significant 1.7-fold decrease for *aur*, 2 for *sspA*, and 1.8 for *spl* transcript levels (**Fig. 6F**), alongside no changes for *scpA* transcription. Lastly, for *sarU* we observed a significant 2.3-fold decrease for *sspA* and 1.7-fold for *aur* transcript levels (**Fig. 6G**), whilst no changes were noted for *scpA* and *spl* transcription. Collectively, almost all of the regulators solely activate protease transcription, with the exception of XdrA, which differentially regulates protease loci in opposing fashions, akin to that observed with Rot and SarS.

**Figure 6.**
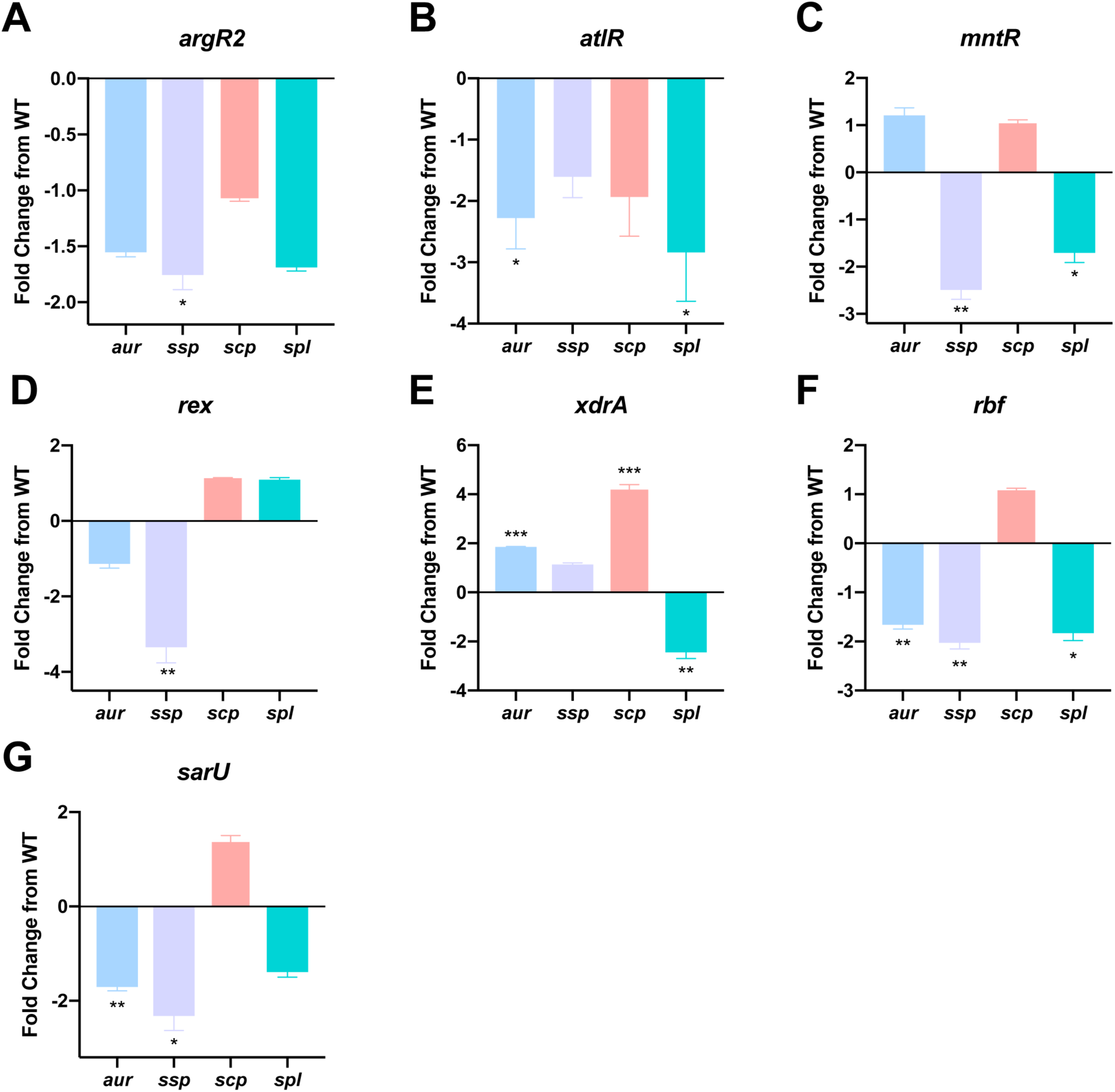
Differential Control of Individual Protease Loci by a Secondary Network of Regulatory Factors. qRT-qPCR was performed to determine transcript levels for *aur, ssp, scp, and spl* in the regulator mutants. The strains used were: WT = USA300 HOU, alongside mutants of *argR2, atlR, mntR, rex, xdrA, rbf,* and *sarU*. RNA was isolated from three independent cultures. The 16s rRNA gene was used as an internal control. Fold change from WT was determined using the 2^−ΔΔC^_T_ method. A Student’s t-test was used to determine statistical significance, *=p< 0.05, **=p< 0.01, ***=p<0.001. Error bars are shown ± SEM.

### Determining the Pathway of Control for the Novel Protease Regulators

In the work above, we identified 14 new regulatory pathways for secreted protease transcription. This data allows us to construct a map of protease regulation for these factors, detailing specific effects on individual protease loci (**Fig. 7**). To delineate the pathway by which these regulators exert their effects, we next assessed their impact on the primary regulators of protease expression considered previously. As such, qRT-PCR analysis was performed on the seven novel protease regulator mutants for *sarA*, *codY*, *rot*, *sarS*, *saeR*, *mgrA*, and *sarR*. SarA, SarR, MgrA, and CodY are able to regulate protease production by direct action, but can also act via control of the Agr quorum sensing system [26, 72–78]. Agr in turn activates secreted protease production during post-exponential phase by inhibiting translation of the negative regulator Rot [79–81]. As such, for completeness, we also included analysis of the *agr* operon in these studies.

**Figure 7.**
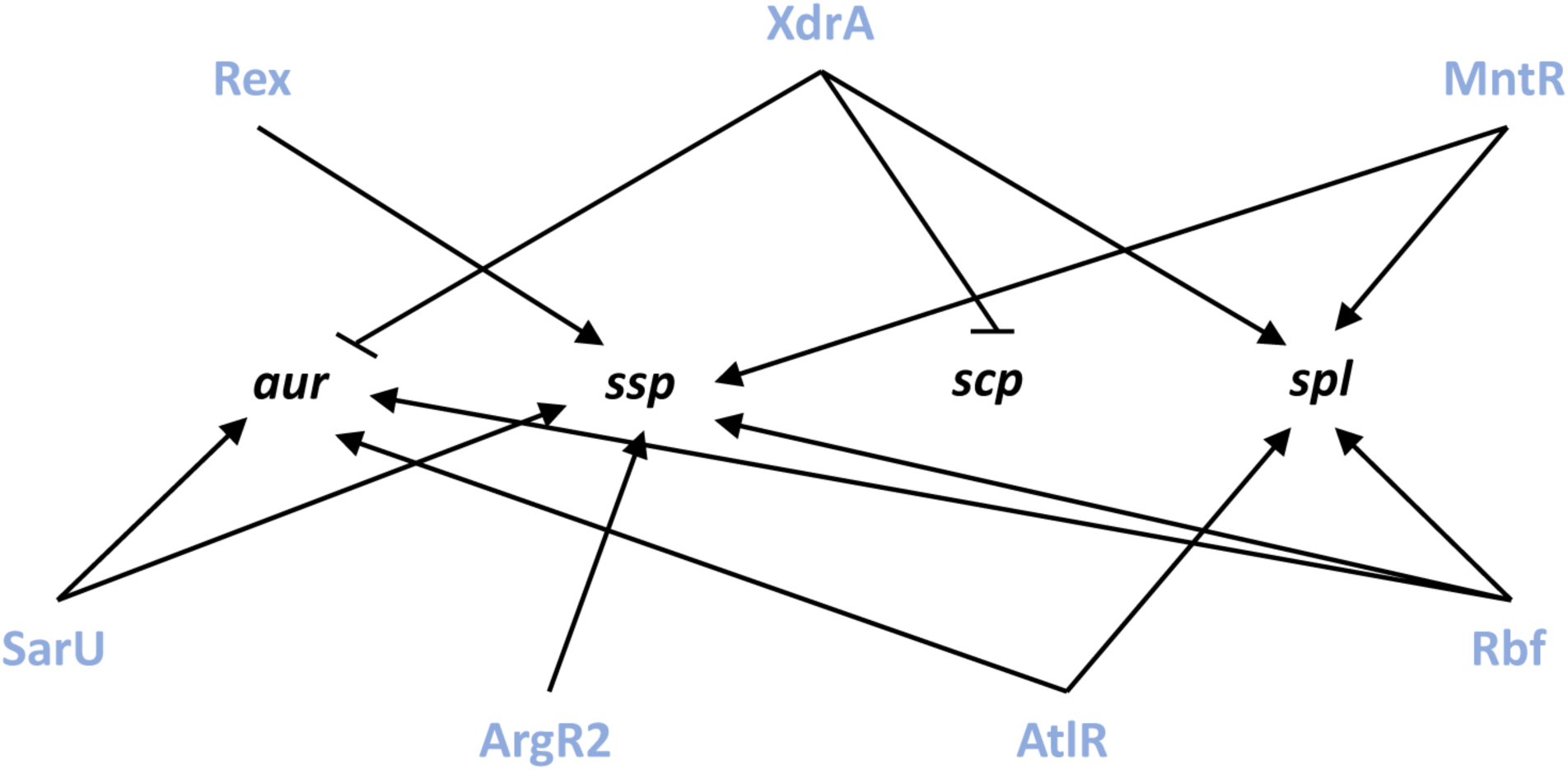
A Novel Regulatory Network Controlling Expression of Extracellular Proteases. Shown are transcriptional regulation events for the seven novel protease regulators on the four individual protease loci. Bars indicate repression and arrows activation.

When data for the *argR2* mutant was analyzed we found no significant changes in expression for any of the primary protease regulators (**Fig. 8A**). As such the changes in *ssp* transcript levels in the *argR2* mutant are either the result of direct action by ArgR2 or are mediated by an as yet unknown circuit. When assessing the *atlR* mutant, a significant 1.4-fold decrease in *saeR* and a 1.5-fold increase in *sarS* transcripts was observed (**Fig. 8B**). The decrease in *saeR* could explain the observed decrease in *spl* expression, as SaeR was shown by ourselves and others to activate *spl* transcription [24, 45]. In addition, the increase in *sarS* expression could explain the decrease in both *aur* and *spl* transcripts, as SarS was shown in this study to repress transcription of *spl* and was shown here and elsewhere to repress *aur* expression [25].

**Figure 8.**
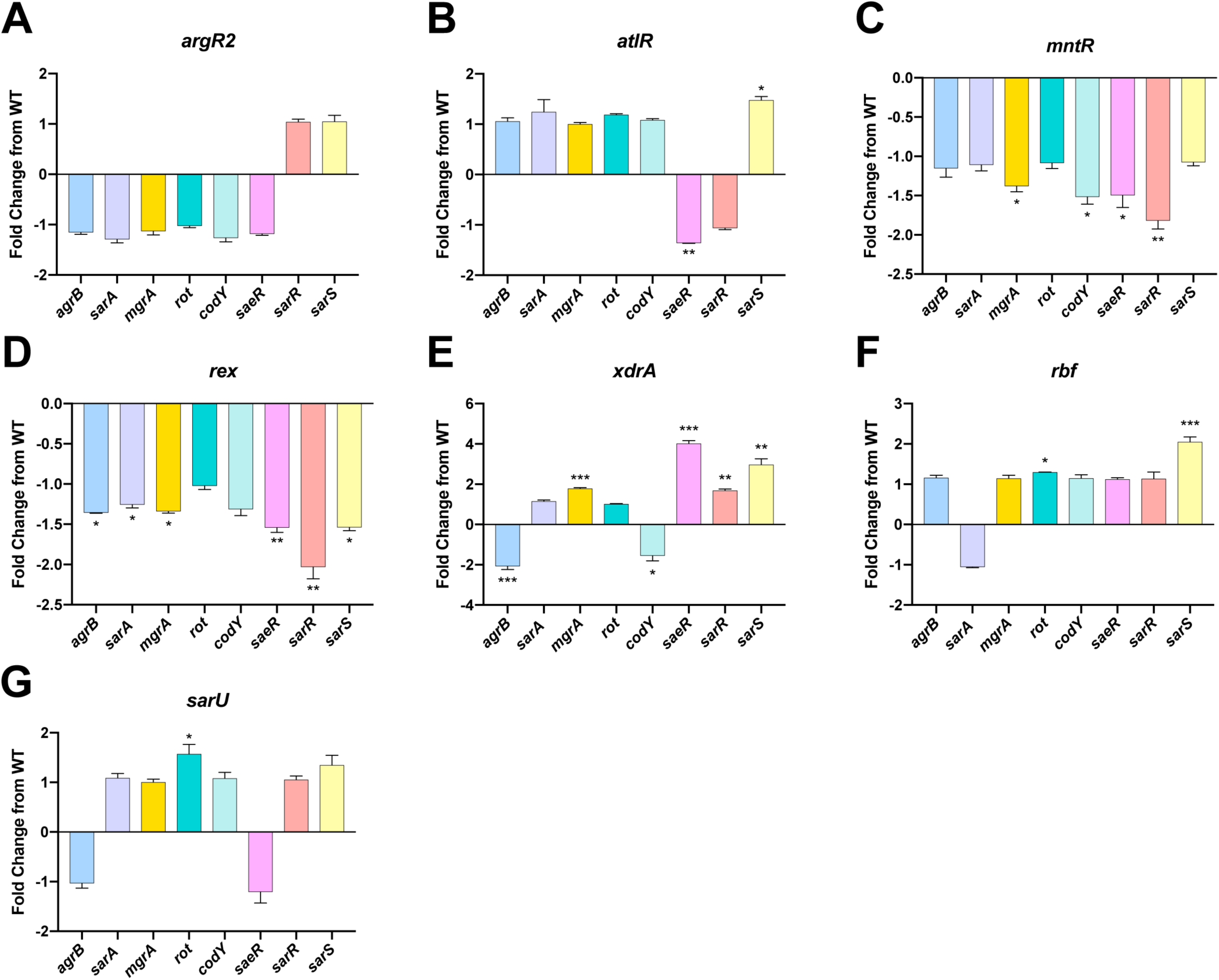
Determining the Pathway of Control for the Novel Protease Regulators. qRT-PCR was performed to determine transcript levels for *agrB, sarA, mgrA, rot, codY, saeR, sarR,* and *sarS* in the regulator mutants. The strains used were: WT = USA300 HOU, alongside mutants of *argR2, atlR, mntR, rex, xdrA, rbf,* and *sarU*. RNA was isolated from three independent cultures. The 16s rRNA gene was used as an internal control. Fold change from the WT was determined using the 2^−ΔΔC^_T_ method. A Student’s t-test was used to determine statistical significance, *=p< 0.05, **=p< 0.01, ***=p<0.001. Error bars are shown ± SEM.

Next, with *mntR* we observed a significant 1.4-fold decrease in *mgrA*, 1.5-fold in *codY*, 1.5-fold in *saeR*, and 1.8-fold in *sarR* transcript levels (**Fig. 8C**). With regards to the decrease in *ssp* and *spl* transcripts, these changes cannot be explained by the decrease in transcription for *codY* as we show that CodY represses both of these loci. The decrease in *saeR* transcript, however, could result in a decrease in *spl* transcription as it has been shown by ourselves and others to be an activator of this operon [24, 45]. Furthermore, the decrease in *mgrA* and *sarR* transcripts could lead to a decrease in *ssp* and *spl* expression, as we confirm the work of others demonstrating that MgrA activates expression for both proteases [25, 34], whilst at the same time newly identifying SarR as acting in a similar fashion.

When exploring the influence of Rex, we observed a significant 1.4-fold decrease in *agrB*, 1.3-fold in *sarA*, 1.3-fold in *mgrA*, 1.5-fold in *saeR*, 2-fold in *sarR*, and 1.5-fold in *sarS* transcript levels (**Fig. 8D**). The changes in *sarA, saeR,* and *sarS* cannot explain the decrease we observed for the *ssp* transcript because, as shown by ourselves and others, both are repressors of *ssp* [9, 10, 25]. However, as we and others have shown that MgrA, SarR, and Agr are activators of *ssp* transcription [9, 21, 25], decreases in their expression could explain our data. When assessing *xdrA*, a significant 2.1-fold decrease in *agrB* and 1.5-fold in *codY* transcript levels were observed (**Fig. 8E**). Additionally, a significant 1.8-fold increase in *mgrA*, 4-fold in *saeR*, 1.7-fold in *sarR*, and 3-fold in *sarS* transcripts was observed. The increase in *mgrA* transcript could explain the increase in *aur* expression as MgrA has been shown here and by others to activate its transcription [25, 34]. Next, as we showed SaeR, SarR, and SarS to be activators of *scp* expression, increases in the transcription of each could result in enhanced *scp* transcript abundance. Additionally, the decrease in *codY* expression could explain the increase in transcript for *aur* and *scp*, as we showed CodY to be a repressor of both. Lastly, the decrease in *spl* transcript levels in the *xdrA* mutant could be explained by either the increase in *sarS* or by the decrease in *agrB* expression, as we show that SarS is a repressor of this loci, whilst it is well known that Agr is an activator of *spl* transcription [10]. Next, with *rbf* we observed a significant 1.3-fold increase in *rot* transcription as well as a 2.1-fold increase for *sarS* (**Fig. 8F**). The decrease in *aur* and *ssp* transcript levels observed in the *rbf* mutant could be explained by the increase in *sarS* expression, as it was shown by ourselves and others to be a repressor for both loci [25]. Further, we show SarS is a repressor of *spl* and as such, the increase in *sarS* could have resulted in the decrease in *spl* transcript. In addition, Rot was shown herein, and by others, to be a repressor for *aur* and *ssp*, and therefore the increase in *rot* transcription could result in the decrease of *aur* and *ssp* expression [23]. Lastly, with *sarU* we observed a significant 1.6-fold increase in *rot* transcription (**Fig. 8G**). In the *sarU* mutant the decrease in *aur* and *ssp* transcription could be explained by the increase in *rot* transcription as it has been shown by ourselves and others to be a repressor of both [23].

### Integrating the Novel Secondary Protease Regulators into the Global Picture of Protease Control

Using the findings from this study, along with existing knowledge, we put forward a comprehensive map of secreted protease regulation (**Fig. 9**). With this knowledge, we are able to identify specific regulatory pathways connecting our novel protease effectors with the major protease regulators. Specifically, with regard to Rbf, it is possible that its repressive effect on *sarS* transcription is through Rot as it was previously shown to activate *sarS* transcription [41, 77], and *rot* transcription is increased in the *rbf* mutant. Next, with MntR, its positive effect on *sarR* transcription are likely occurring through MgrA as it was previously shown that MgrA activates *sarR* transcription [34], and *mgrA* expression is decreased in the *mntR* mutant. As for Rex, its activation of *sarR* transcription could be occurring through MgrA as it has been shown that MgrA activates *sarR* [41, 77], and *mgrA* transcription is decreased in the absence of *rex*. Lastly, with XrdA, it is possible that its represses *saeR* via CodY as its been shown that CodY represses *saeR* transcription [82, 83], and *codY* transcription is decreased in the *xdrA* mutant. Additionally, the negative effect of XdrA on *sarR* and *sarS* transcription could be occurring via MgrA as it was previously shown that MgrA activates *sarR* and *sarS* transcription [34, 77] and *mgrA* transcription is increased in the *xdrA* mutant. Finally, the activation of *agr* by XdrA could by occurring via the MgrA-SarR pathway as SarR has been shown to repress *agr* transcription [74], and as already noted, *sarR* transcription is increased in the *xdrA* mutant.

**Figure 9.**
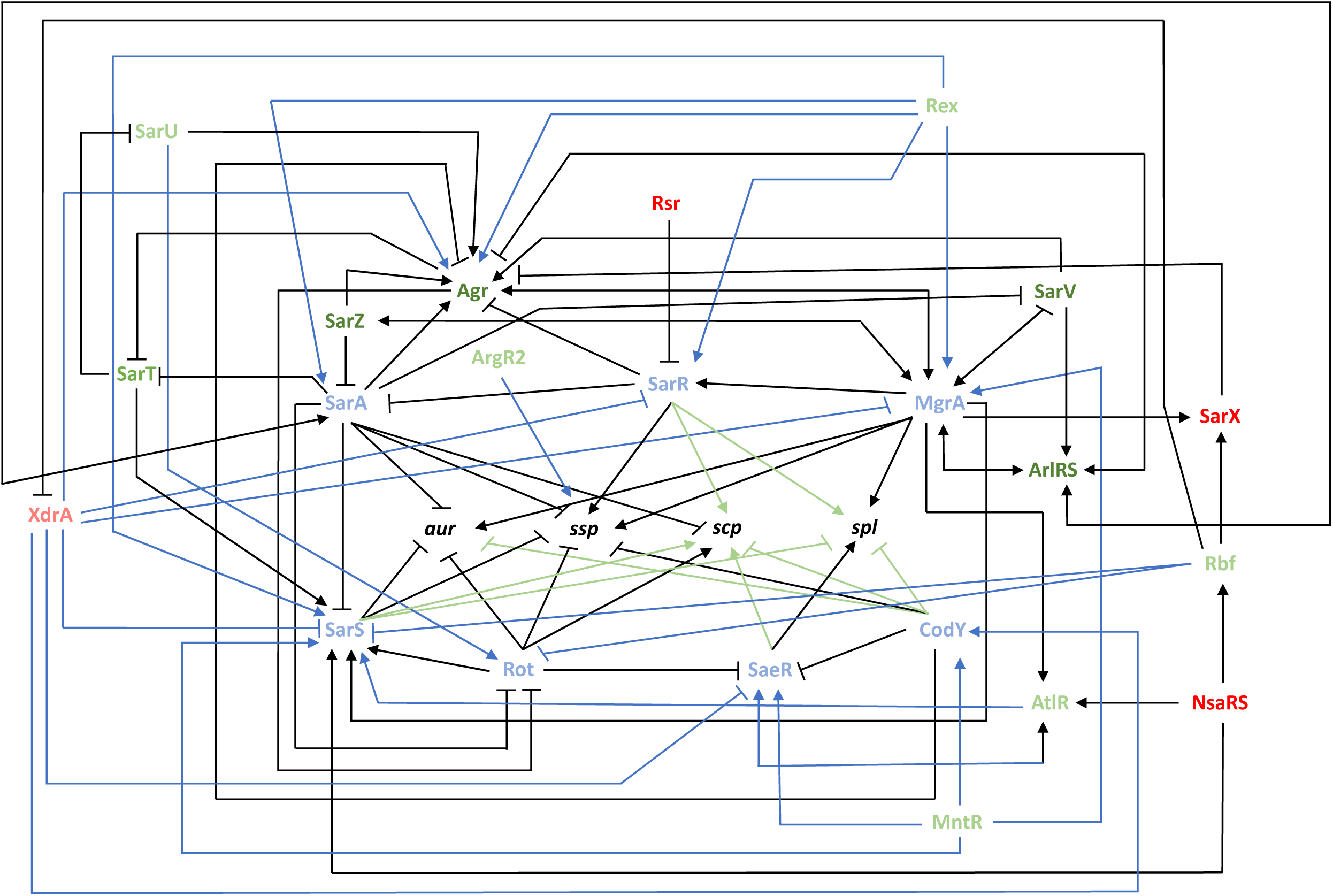
Mapping the Global Network of Extracellular Protease Regulation in *Staphylococcus aureus*. The seven primary regulators of protease expression are shown in blue, whilst factors known to, in turn, regulate their expression, are shown in dark green (activators) or dark red (repressors). The novel regulators identified in this study are shown in light green (activators) or light red (repressors). New regulatory pathways identified herein between the primary regulators and the protease loci are shown in green. New regulatory pathways identified herein between the primary regulators and the novel regulators are shown in blue.

### Concluding Remarks

In this study we set out to completely characterize the loci specific effects of regulatory factors on secreted protease expression. In so doing, we have identified an abundance of novel regulatory nodes controlling their production, and present a comprehensive regulatory circuit that emphasizes the complexity of protease regulation (**Fig. 9**). When one compares this regulatory overview with the literature on virulence factor control in *S. aureus*, it becomes clear that the expansive and complex regulatory circuits that exist to oversee secreted protease expression rivals that of α-toxin and Protein A, which are arguably some of the most important virulence affecting entities produced by this organism [41, 71, 84–91]. Indeed, we suggest that the existence of such a broad network of control speaks to the importance of the secreted proteases to *S. aureus* physiology and pathogenic potential. We also contend that there is a clear and obvious need for such a network, so as to limit or enhance the abundance (and thus activity) of these enzymes. The rationale for this is that a primary function of these enzymes is to control the progression of infection by selectively modulating the stability of individual virulence factors produced by the cell [19]. Thus, in this context, it makes sense that a network of control exists to selectively titrate in or out a given protease (and thus its activity), so as to specifically influence the abundance (or lack thereof) of individual virulence factor(s). This would then facilitate the selective and niche specific pathogenic behaviors of *S. aureus* and provide a basis for control of the broad and varied infection types that is the hallmark of this organism’s disease-causing nature. Further to this, there is abundant evidence in the literature implicating the secreted proteases as facilitating the infectious process by attacking the host and cleaving host proteins. It is thus in line with the above hypothesis that tightly controlling protease activity, by selectively limiting or enhancing their activity in specific niches, is to the advantage of *S. aureus* and its highly effective and efficient infectious process.

## Supplemental Figure Legends

**Supplemental Figure 1. Protease Activity Profiling of Novel Regulator Mutants During Growth.** Gelatin zymography was performed on USA300 HOU WT and mutant strain culture supernatants obtained at the times specified. Culture supernatants were concentrated and ran on a SDS-PAGE gel containing 0.1% gelatin. Strains used are indicated on each gel.

**Supplemental Figure 2. Growth Analysis of Novel Protease Regulator Mutants.** USA300 HOU WT and regulator mutants of: *argR2*, *mntR*, *atlR*, *rbf*, *sarU*, *xdrA*, and *rex* were grown under standard conditions in TSB. Data is from three biological replicates with error bars shown ±SD.

